# Predator-induced selection on urchin activity level depends on urchin body size

**DOI:** 10.1101/655977

**Authors:** Justin Pretorius, James L.L. Lichtenstein, Erika J. Eliason, Adrian C. Stier, Jonathan N. Pruitt

## Abstract

Temporally consistent individual differences in behavior impact many ecological processes. We simultaneously examined the effects of individual variation in prey activity level, covering behavior, and body size on prey survival with predators using an urchin-lobster system. Specifically, we tested the hypothesis that slow-moving purple sea urchins (*Strongylocentrotus purpuratus*) and urchins who deploy extensive substrate (pebbles and stones) covering behavior will out-survive active urchins that deploy little to no covering behavior when pitted against a predator, the California spiny lobster (*Panulirus interruptus*). We evaluated this hypothesis by first confirming whether individual urchins exhibit temporally consistent differences in activity level and covering behavior, which they did. Next, we placed groups of four urchins in mesocosms with single lobster and monitored urchin survival for 108 hours. High activity level was negatively associated with survival, whereas urchin size and covering behavior independently did not influence survival. The negative effect of urchin activity level on urchin survival was strong for smaller urchins and weaker for large urchins. Taken together, these results suggest that purple urchin activity level and size jointly determine their susceptibility to predation by lobsters. This is potentially of great interest, because predation by recovering lobster populations could alter the stability of kelp forests by culling specific phenotypes, like foraging phenotypes, from urchin populations.

## 1 INRODUCTION

Intraspecific differences in morphology and behavior are predicted to shape the outcome of species interactions. For example, prior studies have shown that larger and faster prey species are more likely to escape predation (Einfalt & Wahl, 1997; Hoyle & Keast, 1987; Werner & Hall, 1974). Likewise, individual variation in predator jaw morphology can shape the effect of predators on community composition at both the producer and primary consumer levels (Post, Palkovacs, Schielke, & Dodson, 2008). Parallel lines of evidence suggest that among individual variation in key traits of both predators and primary producers can affect ecosystem structure and function (Bolnick et al., 2011; Bolnick et al., 2003; Wolf & Weissing, 2012). For instance, genetic diversity within primary producer species is correlated with consumer diversity and community resilience, seemingly by increasing the diversity of interaction types (Crutsinger et al., 2006; Hughes & Stachowicz, 2004; Johnson, Lajeunesse, & Agrawal, 2006; Reusch, Ehlers, Hämmerli, & Worm, 2005; Rudman et al., 2015). Understanding how intraspecific variation in morphology and behavior shape the outcome of predator-prey interactions is of specific interest, because these interactions can change consumer abundance and behavior, and therefore, alter the role of predation in driving the stability, function, and resilience of ecosystems (Brown, Laundré, & Gurung, 1999; Miller, Ament, & Schmitz, 2014; Preisser & Bolnick, 2008).

The past few decades have seen a rise in the number of studies focused on intraspecific behavioral variation (Bell, Hankison, & Laskowski, 2009; Sih, Cote, Evans, Fogarty, & Pruitt, 2012). In particular, much literature has examined the effects of temporally consistent individual differences in behavior, or so called “animal personalities” (Carter, Feeney, Marshall, Cowlishaw, & Heinsohn, 2013; Gosling, 2001) and their effect on species interactions (Chang, Teo, Norma-Rashid, & Li, 2017; Lichtenstein, Wright, McEwen, Pinter-Wollman, & Pruitt, 2017; Nannini, Parkos III, & Wahl, 2012). Animal personalities imply that there exists a level of constraint to the behavioral plasticity individuals display in any given context (Dall, Houston, & McNamara, 2004; Dingemanse, Kazem, Réale, & Wright, 2010; Dochtermann & Dingemanse, 2013), meaning that individuals may exhibit behavioral consistency through time and across contexts, occasionally to their detriment (Jones & Godin, 2010; Pearish, Hostert, & Bell, 2013; Pruitt, Riechert, & Jones, 2008). For instance, some prey animals ceaselessly seek shelter regardless of the contextual risk, which can result in decreased opportunities to forage or mate, while other animals may continuously forage regardless of the level of predation risk (Dall et al., 2004; Wilson, Coleman, Clark, & Biederman, 1993). Yet, the available empirical evidence suggests that the effects of prey behavior on the outcome of predator-prey interactions varies considerably among systems (Belgrad & Griffen, 2016; Keiser, Ingley, Toscano, Scharf, & Pruitt, 2017), and even varies based on the traits of the individuals (predator and prey) involved (DiRienzo, Pruitt, & Hedrick, 2013; McGhee, Pintor, Suhr, & Bell, 2012; Sweeney et al., 2013).

Here, we use the purple urchin (*Strongylocentrotus purpuratus*), and the California spiny lobster (*Panulirus interruptus*), to evaluate the effect of prey behavioral tendencies and size on prey survival. Urchins are ecologically notable because of their ability to consume large quantities of giant kelp (*Macrocystis pyrifera*) and therefore alter the state of coastal kelp communities (Estes & Duggins, 1995; Harrold & Reed, 1985; Pearse, 2006). Across the wide geographic distribution of purple urchins along the west coast of North America (Ebert, 2010), urchins are eaten by a diversity of predatory species including the sunflower sea star (*Pycnopodia helianthoides*), otters (*Enhydra lutris*), sheephead (*Semicossyphus pulcher*), and California spiny lobsters (*Panulirus interruptus*). These predators can play key roles in regulating herbivory by urchins on giant kelp (Burt et al., 2018; Caselle, Davis, & Marks, 2018). However, the role of variation in urchin size structure and individual behavioral variation in their susceptibility to predation remains mysterious, even though such variation in mortality could explain variation in community susceptibility to urchin-driven state shifts, like between kelp forests vs. urchin barrens, a process often referred to as a “tipping point” (Pruitt et al., 2018; Selkoe et al., 2015). We focused on how two behavioral traits in might determine prey survival in the presence of spiny lobsters: propensity to move (activity level), which can increase or decrease encounter rates with predators (Huey & Pianka, 1981; Skelly, 1994), and urchin’s tendency to conceal themselves with substrate (covering behavior). Although the function of urchin covering behavior appears to vary among species (i.e., protection from wave action; Dumont, Drolet, Deschênes, & Himmelman, 2007), we predicted that urchins would be safer from predators while buried in substrate.

We tested the hypothesis that slow-moving purple urchins and urchins with a greater tendency to cover themselves and/or larger urchins will enjoy greater survival during staged encounters with sit-and-pursue predators. The former is predicted by Locomotor Crossover Hypothesis proposed by Huey and Pianka (1981), which posits that predators should tend to encounter and consume prey of the opposing activity level (active predators consume sedentary prey, whereas sit-and-pursue predators consume active prey). To probe the role of size, we tested whether urchin size increases urchin survival jointly with urchin behavioral traits. We further evaluated whether prey behavior measured in isolation is correlated with prey behavior during real encounters with predators. Consistency across testing conditions is a commonly untested assumption in animal personality studies, usually overlooked because of logistical limitations. However, the conspicuousness of the urchin covering behavior and the slow movements of predator and prey in this study made ongoing assessment of this behavior achievable. Together, this system allowed us to evaluate how the behavioral and morphological traits of a prey determine their susceptibility to predation.

## 2 MATERIALS AND METHODS

### 2.1 Animal collection and maintenance

Fifty *Strongylocentrotus purpuratus* were opportunistically collected by hand from the Goleta sewer pipe (34° 24.851 N; 119° 49.749 W) and 120 others were collected from Goleta Pier (34° 24.847 N; 119° 49.718 W, Goleta, CA, U.S.A.) in the summer of 2017. A total of 170 *S. purpuratus* were used for the experiment. The three lobsters (*Panulirus interruptus*) used in the experiment were transferred into our possession from another research facility on the campus of the University of California at Santa Barbara (UCSB) on 16 August 2017. The three *P. interruptus* used in the study were female, 10.5-11cm in carapace length, and were collected by hand from the Santa Barbara Channel from May to June 2017.

All *S. purpuratus* were provided brown algae (*Nereocystis luetkeana*) *ad libitum* every other day from 01 July 2017 to 01 November 2017. Purple urchins were housed in four separate 90.0L cm × 141.0W cm × 40.5H cm seawater (9-11° C) flow through A-frame tanks until their behavioral assays. During this time, we measured the total diameter (test and spines) of each urchin using digital calipers. They had average diameter of 7.7 ± 0.20 SE cm and a range of 4.5-11.5 cm. After purple urchins had been assayed for covering behavior, individuals were transferred into numerically labeled 17.15L cm × 11.90W cm × 7.00H cm plastic containers in the same seawater flow through tanks. Plastic containers housing individual purple urchins had fourteen holes of 16.0 mm diameter drilled into their sides to allow seawater flow through. *P. interruptus* were housed in similar seawater flow-through A-frame tanks bisected by plastic dividers with twelve 16mm holes, creating two identical lobster enclosures (90.0L cm × 70.5W cm × 40.1H cm). Each lobster enclosure contained three cinder blocks, two of which supported the plastic dividers. All animals were exposed to open air conditions and natural day-night cycles for the duration of the experiment. This experiment was conducted at the UCSB Marine Laboratory from June to November 2017.

### 2.2 Repeatability overview

To estimate short-term behavioral repeatability, we ran ten urchins though 6 trials assessing movement velocity (activity) and covering percentage (covering behavior) over the span of six days. We ran the urchins through two covering behavior tests separated by an hour, and the next day, we ran the urchins through two activity level tests separated by an hour. We repeated this two-day cycle for six days. Therefore, we assessed the repeatability of activity level and covering behavior with six iterations across three testing days for each assay type in ten focal individuals. These ten testing urchins were not used in predator-prey trials.

The purple urchins not selected for repeatability estimates underwent only two trials of each type, conducted under identical conditions (n_individuals_ = 160) separated by an hour. The average value of the two activity and covering behavior measurements for each urchin served as our activity and covering behavior predictor variables in our statistical analyses predicting prey survival.

### 2.3 Measuring urchin activity level

To determine urchin activity level, we estimated their average velocity in open field arenas. The arena was a 40.60L cm × 32.40W cm × 15.25H cm plastic container fashioned with 6 holes of 16.0 mm diameter drilled into its sides and submerged in flow through seawater. The container was completely submerged in seawater within a chilled seawater flow through plastic A-frame tank. We placed a 28.0 cm × 30.5 cm particleboard with a 4 cm grid within each activity container to track urchin movement distance. The board was covered with transparent saran wrap to help the urchins cling to the board. We used movements between squares to calculate an average movement velocity (mm/s) for each individual, one urchin at a time, across a ten-minute trial. We deemed urchins with higher velocities to be more active. This test is very similar to an open field test, a classic metric of activity level (Carter et al., 2013). We performed two such activity trials on a single testing day separated by an hour on each of 160 urchins. We averaged the two scores to obtain a single estimate of each urchin’s activity level in a novel environment. Also, two activity containers were run at a time.

### 2.4 Measuring sea urchin covering behavior

To estimate urchins’ shelter seeking tendencies, we measured their propensity to cover themselves with pebble substrate. Covering behavior resembles common metrics of “boldness” in the personality literature and has antipredator benefits in some species of urchins (Amsler, McClintock, & Baker, 1999). However, the behavior can be cued by wave action and sunlight in other urchin species (Dumont et al., 2007), although urchins deploy these behaviors in the absence of both (Pawson & Pawson, 2013). We examine here whether covering behavior might provide an antipredator benefit to purple urchins with predators during staged encounters.

Individual purple urchins (n = 170) were haphazardly placed into pairs and then tested under social conditions. Pairs of asymmetrically sized urchins (difference in test [spherical] diameter of 10.0 mm or greater) were created to track individual identity during these trials. Urchins were tested in pairs because of the time constraints of testing so many urchins. Pairs of purple urchins were placed into 40.60L cm × 32.40W cm × 15.25H cm plastic containers with transparent lids fashioned with 6 holes of 16.0 mm diameter drilled into their sides to allow chilled seawater to flow through. These were the same containers used for the activity level test, but for covering behavior trials, the bottoms of these containers were layered with 2.0-3.0 cm of fresh composite pebbles of 0.3-1.5 cm diameter, rather than a particle board grid. These containers were then completely submerged in chilled seawater flow within plastic A-frame tanks entirely separate from their home A-frame tank.

Once the pair of purple urchins were placed into the container and submerged, they were permitted one hour to move about their arenas and cover themselves. After this time, we opened the arena and estimated the percent cover (test + spines) of each urchin by eye to the nearest 5%. These estimates were taken by a single observer (Pretorius J). We performed two trials in a testing day separated by an hour. We then averaged each urchin’s scores to obtain a single estimate of urchins’ covering behavior.

### 2.5 Spiny lobster vs. sea urchin staged interactions

Each staged predator-prey interaction consisted of a cohort of 4 haphazardly selected urchins and a single predator lobster. Lobsters were maintained under identical feeding conditions (*ad libitum* mussels interspersed with purple urchins to ensure they did eat urchins) and then starved for four days prior to the initiation of our trials, and all purple urchins had already been assayed for activity and covering behavior. Because we had three lobsters, we ran three of these trials at once. Predators were permitted to interact with prey for 108 hours during our trials, because this duration resulted in an urchin mortality rate over 50%.

Staged interactions were conducted in natural seawater (9-11° C) flow-through tanks divided by plastic barriers perforated by twelve 16 cm holes. The bottom of each arena (90.0L cm × 40.5W cm × × 70.5H cm) was layered with 2.5-4.0 cm of pebbles of diameter 0.3-1.5 cm and contained three submerged cinderblocks to permit prey to hide from predators in physical retreats or, most commonly, within the substrate. The sides of the tank likewise served as physical refuge for prey, allowing them to climb up the sides of the container and out of reach of the lobsters.

To distinguish individual purple urchins throughout staged interactions, one of four colors (yellow, green, pink, or blue) of acrylic nail polish was lightly dotted on the side of a few of each individual’s peripheral spines. Once acrylic nail polish had been applied to every purple urchin within a group of four, all four purple urchins were simultaneously placed by hand into four corners (quadrants) of each arena containing a lobster. Care was taken to place urchins outside of the lobster’s field of view. For each occasion, staged interactions began at 14:30 hours. We recorded each urchin’s covering percentage and the quadrant they occupied every hour for the first 24 hours and every six hours for the next 84 hours. We estimated activity during trials by using movement between quadrants. After 108-hours, a thorough search of every mesocosm was conducted. We recorded which urchins had survived by their color markings. We repeated these trials four times separated by a week using entirely new urchins, so each lobster was used in four trials. The flow-through A-frame tanks for predation trials were housed on a separate concrete patio surrounded by chain link caging to prevent mammalian predators (raccoons, etc.) from eating captive marine fauna, which is required for the vertebrates housed in other nearby enclosures. Thus, urchins that went missing during our trials had few escape options and no other likely predators.

### 2.6 Statistical Methods

#### 2.6.1 Repeatability of behavior

The repeatability of each behavioral trait was determined using the rptR package (Stoffel, Nakagawa, & Schielzeth, 2017) in R version 3.4.3 (R Development Core Team, 2010). We used this package to fit mixed models with ‘individual ID’ and “testing day” (1-3) as random effects, “trial number” (1/2) as a random effect nested within testing day, and each behavioral test was set as the response variable. These models estimate repeatability as the proportion of variance determined by ‘individual ID” to the total behavioral variance. The rptR package constructs 95% CI of the repeatability estimates using 1000 bootstrap iterations of these repeatability estimations. The activity level repeatability models were linear mixed model, because a Gaussian distribution best fit the data. The covering behavior repeatability models were generalized linear mixed effects models fit with a Poisson distribution, because a Poisson distribution best fit the data. Repeatability estimates are deemed significant if their 95% confidence interval do not overlap 0.

#### 2.6.2 Spiny lobster-sea urchin staged interactions

To investigate the factors predicting urchin survival, we deployed a generalized linear mixed model (GLMM) fit with a binomial distribution in the lme4 package in R (Bates, Mächler, Bolker, & Walker, 2014). Our full GLMM had (0/1) urchin survival as its response variable, and its predictor variables were urchin average activity level, average urchin covering behavior, urchin diameter, and all of their interaction terms. Lobster ID and trial ID were included as random effects in our analysis. To build the model with the greatest likelihood, we ran this full model through backwards stepwise BIC model selection (Chambers & Hastie, 1992).

#### 2.6.3 Cross contextual covering behavior and activity

We tested (1) whether urchins’ pre-mesocosm average covering percentage predicted covering behavior in the staged interactions, and (2) whether pre-mesocosm average velocity (activity level) correlated with the frequency of movement between quadrants of staged interactions. However, urchins varied in their number of behavioral observations collected during staged interactions, because urchins varied in how long they survived. Therefore, we weighted urchins’ average covering percentage and frequency of movement between quadrants during staged interactions by probability of survival estimated. Survival probability was estimated using a binomial logistic regression with survival as the response variable and activity level, covering behavior, diameter, activity by covering behavior, activity by diameter, and covering behavior by diameter interaction terms as explanatory variables. To calculate weighted average covering percentage and frequency of movement between quadrants, we multiplied centered individuals’ average covering percentage and frequency of movement by individuals’ probability of survival respectively. Weighted average covering percentage and weighted frequency of movement between quadrants were non-Gaussian. Therefore, we used Spearman’s rank-order tests to assess the correlation between pre-interaction covering percentage and weighted in situ covering percentage, and the correlation between pre-interaction velocity and weighted in situ movement between quadrants.

## 3 RESULTS

### 3.1 Repeatability of behavior

Urchins behaved consistently across time in their average velocity/activity level (Repeatability [95% CIs] = 0.30 [0.03, 0.57]) and average covering percentage (Repeatability [95% CIs] = 0.73 [0.38, 0.87]). Between-individual differences in covering behavior are notably very high in this test population (Bell et al., 2009). Both of these behaviors are repeatable across a period longer than the lobster predation trials.

### 3.2 Spiny lobster-sea urchin staged interactions

Urchins had a survival rate of 33.33 ± 6.88 % SE. The best model contained urchin dimeter, activity level, and covering behavior (BIC = 75.3, ΔBIC = 3.5). Slow-moving purple urchins survived more frequently than fast-moving urchins (Figure 1; Activity Level: ß = −2.99 ± 1.30 SE, z = −2.30, p = 0.02). However, the effect (Figure 2A; Activity Level × Diameter: ß = 0.16 ± 0.08 SE, z = 1.97, p < 0.05). Urchin covering behavior, on its own, had no direct effect on urchin survival (Covering behavior: ß = −0.01 ± 2.37 SE, z = −0.04, p = 0.98). However, smaller urchins that exhibited greater covering behavior were more likely to survive, whereas urchin covering behavior appeared unrelated to survival among large urchins (Figure 2B; Covering behavior × Diameter: ß = −0.38 ± 0.18 SE, z = −2.12, p = 0.03). Larger urchins appeared to be slightly more likely to survive than small urchins, but not significantly so (Diameter: ß = 0.06 ± 0.03 SE, z = 1.68, p = 0.09), and Lobster ID and trial iteration failed to explain a significant component of the variation in urchin survival, suggesting that survival rates were homogenous across mesocosms.

**FIGURE 1.**
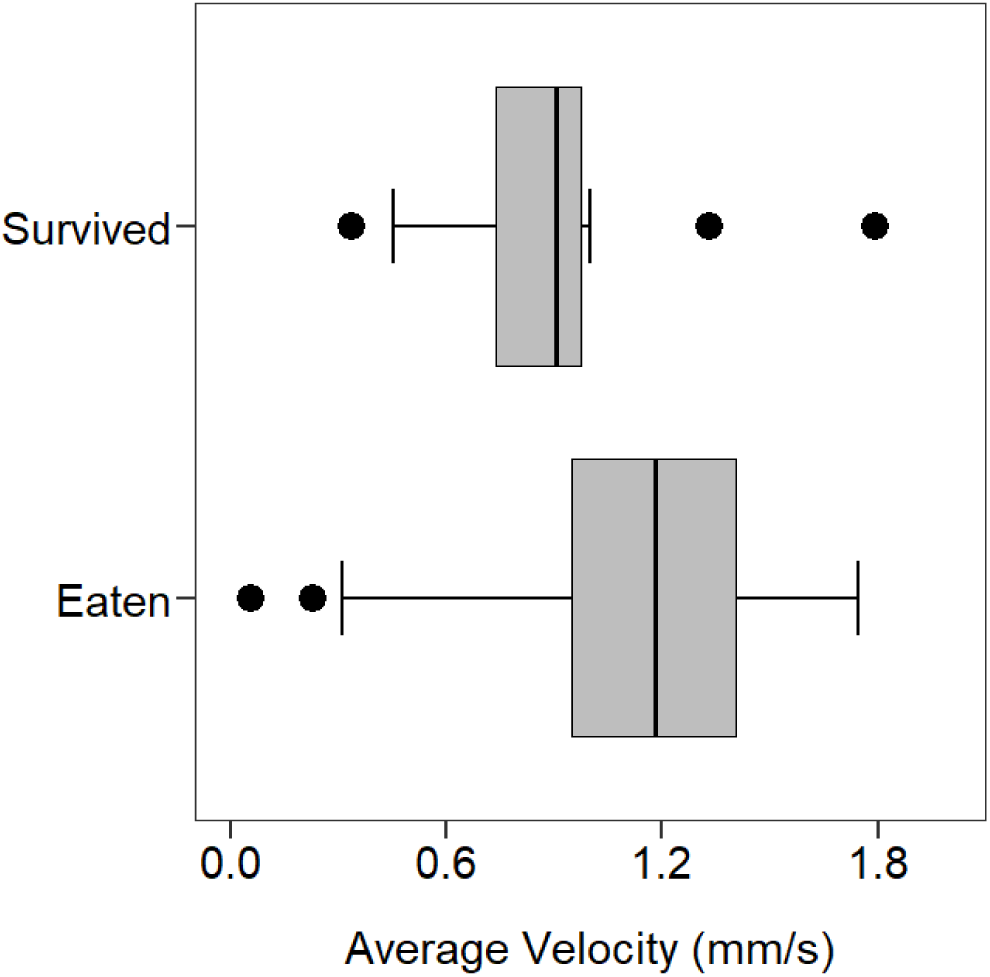
Box plots depicting differences in average velocity between surviving and eaten *S. purpuratus* in staged interactions with *P. interruptus*. Dots denote outliers, whiskers represent 10^th^ and 90^th^ percentiles, gray boxes indicate interquartile range: calculated median inclusive, and the central line denotes the median velocity.

**FIGURE 2.**
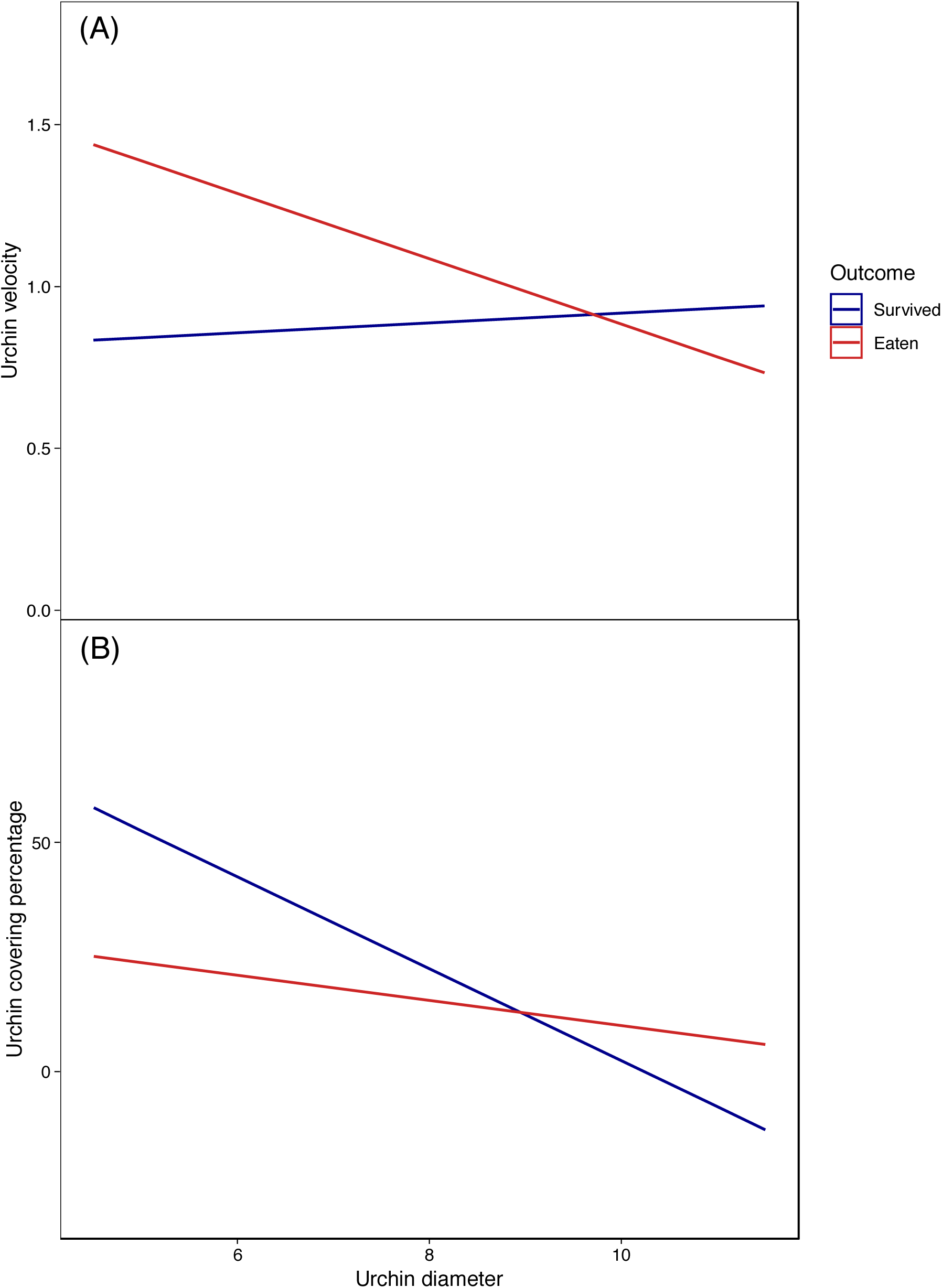
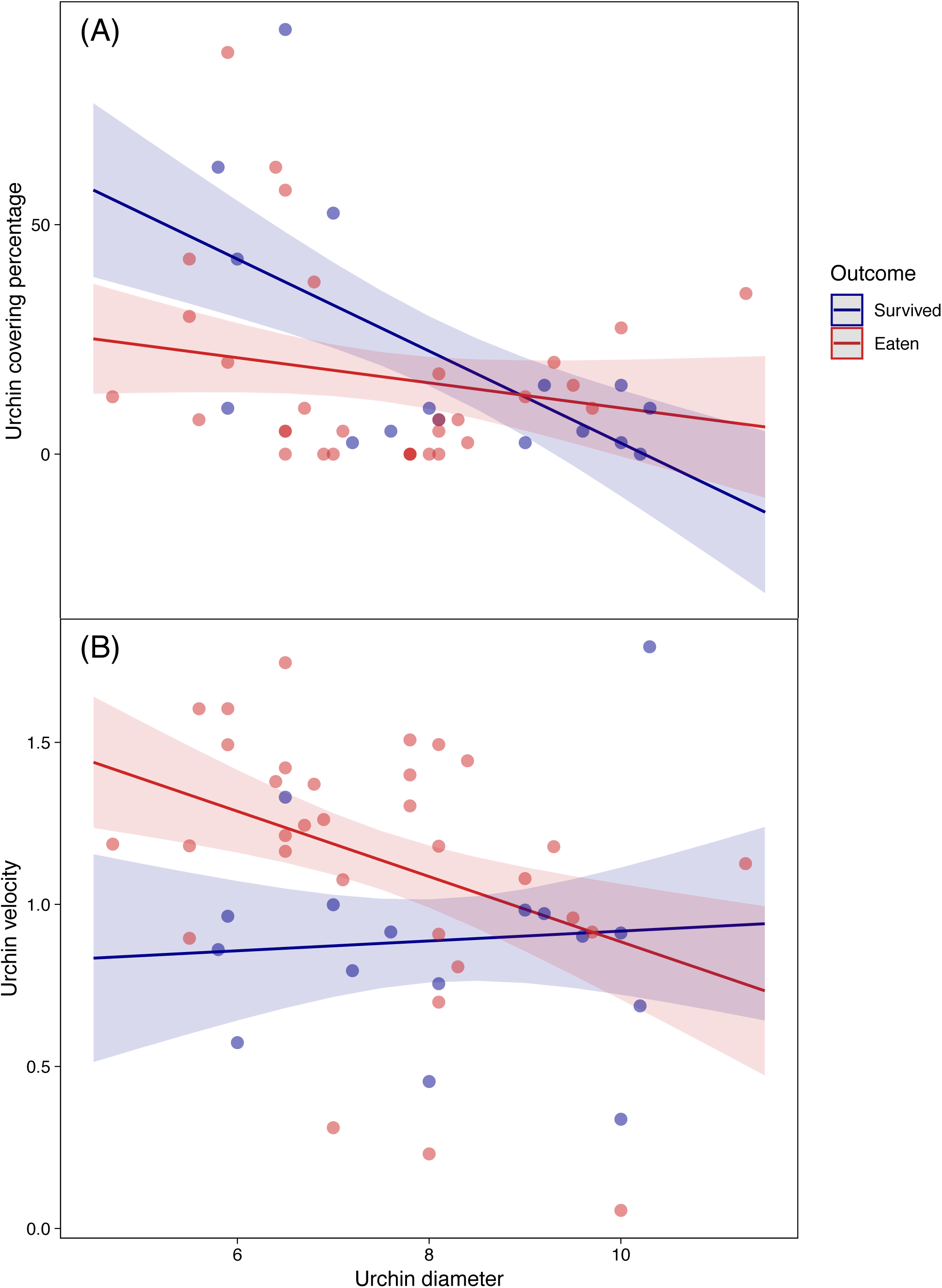
Synergistic effects of covering behavior and size (panel A, top) and activity and size (panel B, bottom) on identity of *S. purpuratus* that survived (white circles) and were eaten (black circles) during staged interactions with *P. interruptus*.

### 3.3 Cross contextual covering behavior and activity

Individual-level differences in average covering percentage were correlated with weighted average covering percentages during staged interactions (Kendall’s tau: Tau = 0.26, p = 0.01). Individual activity, however, seemingly did not persist across contexts, as individual-level differences in average velocity were not correlated with weighted frequency of movement between quadrants during staged interactions (Kendall’s tau: Tau = 0.04, p = 0.69). However, activity level was measured somewhat differently in pre-trial assays during the trial itself, which obscures the interpretability of this result.

## 4 DISCUSSION

Here we evaluated the effects of individual size (test diameter), activity (velocity), and covering behavior on prey survival using California spiny lobsters and purple sea urchins. We found that slow-moving urchins were more likely to survive encounters with lobsters (Figure 1), whereas covering behavior did not predict urchin survival on its own. This is consistent with findings suggesting that active prey are more likely to succumb to predation (Huey & Pianka, 1981; Perry, 2007), particularly with sedentary predators, and that urchin covering behavior does not regularly predict survival in a direct manner (Zhao et al., 2014). Urchin size also did not predict survival directly. Instead, urchin size interacted with urchin activity level and covering behavior to determine urchin survival. Specifically, when purple sea urchins were small, high activity level and low covering behavior reduced their chances of survival, but when urchins were large, their behavior predicted their survival less clearly (Figure 2). Taken together, these findings convey that, even in confined experimental conditions, urchin covering and movement behavior helps to determine survival with predators, but that the ideal behavioral phenotype for survival is contingent on size.

Classical theory on predator-prey interactions predicts that more active animals will be more susceptible to predation (Lima & Dill, 1990). However, more nuanced theoretical frameworks predict that the relationship between prey activity patterns and survival will depend on the foraging mode of the predator (Huey & Pianka, 1981; Perry, 2007; Sweeney et al., 2013). In particular, high activity levels are predicted to be especially costly when prey interact with sit-and-wait or sit-and-pursue predators, which rely on prey movement for encounters. Our results here lend support to both predictions. We found that fast-moving purple urchins were more likely to succumb to lobsters than sedentary urchins. This outcome is consistent with classical optimal foraging and predator-prey interaction theory. However, the fact that spiny lobsters deploy sit-and-pursue foraging tactics (MacDiarmid, Hickey, & Maller, 1991; Tegner & Levin, 1983) suggests that this outcome should be even more likely. The effect of urchin movement speed also deteriorated with urchin size. This finding suggests that activity and covering behavior only mediates survival for smaller and easier to eat urchins. When urchins reach a certain size, they can safely move about with less risk of being eaten (Brooks & Dodson, 1965). It is possible that the detrimental effect of activity level may be driven by the relatively small size of our arenas. Taken together, the age and size structure of urchin populations may determine whether and how lobster predation applies selection pressure on urchin behavior.

Our findings suggest that urchin behavior may potentially play a role in driving the dynamics of kelp forest ecosystems at fine spatial scales. Specifically, smaller (e.g., younger or resource deprived) and more active sea urchin populations may be more susceptible to predation by spiny lobsters, which may alter the role that lobsters play in driving the dynamics of urchin prey. Recent studies have emphasized the importance of size-structured interactions between lobsters and urchins in driving the dynamics of trophically-linked species within the kelp forest (Dunn, Baskett, & Hovel, 2017). However, the role of intraspecific variation in urchin behavior in mediating these size-structured interactions and the resulting effects on kelp forest ecosystem dynamics is not well-studied. A comprehensive integration of intraspecific variation in urchin behavior in driving interactions between urchins and their predators, as well as urchins and kelp, will require a deeper understanding of how urchin behavior shapes their impacts on kelp (amount consumed, foraging mode, etc.) and how conflicting pressures, like the introduction or reintroduction of other predators, including sea otters, or competitors, might generate conflicting selection pressures on prey behavior.

## Conclusions

We found that purple urchin size and behavioral tendencies synergistically determined their susceptibility to predation by spiny lobsters, which are known predators of urchins in nature (Lafferty, 2004; Tegner & Levin, 1983). Understanding the links between prey traits and performance is central to understanding trait evolution and, in the case of this system, potentially important for predicting community level outcomes. Urchin herbivory can defoliate kelp forests, thereby precipitating local loss of biodiversity (Graham, 2004) and altering the value of fisheries (Hamilton, Caselle, Malone, & Carr, 2010; Pearse, 2006). Our results here are a tentative first step towards leveraging the predictive value of intraspecific trait variation to predict how key players in kelp forest ecosystems interact. Moving forward, the challenge is to document how other environmental pressures cause alterations to the trait distributions of predator and prey populations, either in the form of selection or via phenotypic plasticity, and either trophic level’s effects on community level outcomes. We reason that a comprehensive understanding of the role of predator and prey traits in this system will ultimately enable us to better predict or even mitigate urchin generated transitions in kelp forest states. There remains much to be learned, but our results here hint that the effects of individual variation in this system are likely to be strong.

## ACKNOWLEDGEMENTS

We are indebted to the California Coastal Commission for issuing research and collection permits (SCP). We would also like to thank Christoph Pierre for collecting the animals for these studies and assisting in their laboratory maintenance. Funds for this work were generously provided by the University of California, Santa Barbara as start-up to J.N.P., NSF grant awards to J.N.P. (No. 1352705 and No. 1455895), and an NIH grant award to J.N.P. (No. R01GM115509).

